# Alterations of the pulmonary immunity are independent of spontaneous pneumonia in an experimental model of ischemic stroke

**DOI:** 10.1101/618314

**Authors:** Breanne Y. Farris, Kelly L. Monaghan, Courtney D. Amend, Wen Zheng, Heng Hu, James E. Coad, Xuefang Ren, Edwin C.K. Wan

**Affiliations:** Department of Microbiology, Immunology, and Cell Biology, West Virginia University School of Medicine, Morgantown, West Virginia, United States of America; Department of Physiology and Pharmacology, West Virginia University School of Medicine, Morgantown, West Virginia, United States of America; Experimental Stroke Core, Center for Basic and Translational Stroke Research, West Virginia University School of Medicine, Morgantown, West Virginia, United States of America; Pathology Laboratory for Translational Medicine, West Virginia University School of Medicine, Morgantown, West Virginia, United States of America; Department of Neuroscience, West Virginia University School of Medicine, Morgantown, West Virginia, United States of America; Rockefeller Neuroscience Institute, West Virginia University School of Medicine, Morgantown, West Virginia, United States of America

## Abstract

Stroke-associated pneumonia (SAP) is a major cause of mortality in patients who have suffered from severe ischemic stroke. Although multi-factorial in nature, stroke-induced immunosuppression plays a key role in the development of SAP. Previous studies of focal ischemic stroke induction, using a murine model of transient middle cerebral artery occlusion (tMCAO) have shown that severe brain damage results in massive apoptosis and functional defects of lymphocytes in the spleen, thymus, and peripheral blood. However, how immune alternations in remote tissues lead to a greater susceptibility to lung infections is not well-understood. Importantly, how ischemic stroke alters immune-cell fates, and the expression of cytokines and chemokines in the lungs that directly impact pulmonary immunity, has not been characterized. We report here that ischemic stroke increases the percentage of alveolar macrophages, neutrophils, and CD11b^+^ dendritic cells (DCs), but reduces the percentage of CD4^+^ T cells, CD8^+^ T cells, B cells, natural killer (NK) cells, and eosinophils in the lungs. The depletion of immune cells in the lungs is not caused by apoptosis, cell infiltration to the brain, or spontaneous pneumonia following ischemic stroke as previously described, but correlates with a significant reduction in the levels of multiple chemokines in the lungs, including: CCL3, CCL4, CCL5, CCL17, CCL20, CCL22, CXCL5, CXCL9, and CXCL10. These findings suggest that ischemic stroke negatively impacts pulmonary immunity to become more susceptible for SAP development. Further investigation into the mechanisms that control pulmonary immune alternations following ischemic stroke may identify novel diagnostic or therapeutic targets for SAP.

## Introduction

Stroke is the fifth highest cause for overall mortality and the leading cause of long-term disability in the United States [1]. A major cause of mortality is the acquisition of stroke-associated pneumonia (SAP). It was long thought that dysphagia-mediated aspiration following stroke is the sole cause of SAP. However, it is now clear that the stroke-induced immunosuppression is a critical risk factor for acquiring SAP [2], and although it has been recognized for more than 15 years [3], the cellular and molecular mechanisms that trigger this event are still not well-defined. One possibility is that inflammatory immune cells infiltrating into the central nervous system (CNS) from the periphery during re-oxygenation (reperfusion) causes transient exhaustion of immune cells in the periphery, leading to immunosuppression [4]. Another explanation currently under active investigation is that brain damage in stroke triggers the hypothalamic-pituitary-adrenal (HPA) axis and sympathetic nervous system (SNS) in an attempt to dampen further inflammation within the CNS, causing bystander peripheral immunosuppression. These pathways stimulate the secretion of glucocorticoid and catecholamine respectively, both of which are natural immunosuppressive agents. The importance of neurological and endocrinal control of stroke-induced immunosuppression was demonstrated in mouse models with the use of a β-adrenoreceptor blocker propranolol to prevent post-stroke pneumonia [3, 5]. However, a recent historical cohort study concluded that the β-blocker therapy does not reduce the risk of SAP in humans [6]. Therefore, detailed investigation on how different immune cell types are affected quantitatively and functionally after stroke is critical for designing more specific therapeutic strategies.

Current knowledge on the immunological responses following stroke were mostly generated using the murine transient middle cerebral artery occlusion (tMCAO) model of ischemic stroke induction. So far, immune phenotypes that are linked to stroke-induced immunosuppression include 1) induction of apoptosis of lymphocytes (T cells, B cells, and natural killer (NK) cells) in the blood, spleen, and the thymus [3]; 2) reduced expression of proinflammatory TNF but increased expression of immunosuppressive IL-10 in blood and splenic monocytes [7]; 3) reduction of IFN-γ expression in blood T cells and liver natural killer T (NKT) cells [3, 8]; and 4) impaired neutrophil chemotaxis [9]. Among these mechanisms, the reduction of IFN-γ in liver NKT cells seems to be the most relevant, as propranolol treatment in mice following ischemic stroke restores IFN-γ production in these cells [8]. However, currently there is no explanation for how immune cell defects occurring in remote organs, such as the liver and the spleen, lead to bacterial infection in the lungs. Additionally, there is little information on how stroke-induced immunosuppression affects the immune-cell niche and functions in the lungs, which directly impact pulmonary immunity.

Here, we report that the immune-cell niche in the lungs is altered following the tMCAO-mediated ischemic stroke induction, with a significant depletion of lymphocyte populations. Importantly, these changes do not result in spontaneous pneumonia as previously described, but rather they correlate with the reduction of multiple chemokine levels in the lungs, which may represent an immunosuppressive event that accounts for a greater susceptibility to bacterial infection following ischemic stroke.

## Materials and methods

### Animals

Eight to 12-week-old male C57BL/6J mice, weighing 25-30 g were used (The Jackson Laboratory, Bar Harbor, ME). The mice were housed under specific-pathogen-free conditions in the vivarium at West Virginia University Health Sciences Center. Mice were housed according to the Institutional Animal Care and Use Committee (IACUC) guidelines on a 12-hour light/dark cycle and fed and watered *ad libitum*. All protocols and procedures performed were approved by the IACUC of West Virginia University.

### Transient middle cerebral artery occlusion (tMCAO)

Mice were anesthetized with 4-5% isoflurane until a deep plane was reached and animals did not respond to toe-pinch stimulus. Anesthesia was maintained using a nose cone during surgery with 1-2% isoflurane in oxygen enriched air. The common carotid artery and external carotid artery were exposed. Temporary suture was applied to the common carotid artery to stem the flow of blood for filament placement, and an incision was made in the external carotid artery. tMCAO was induced by inserting a monofilament into the external carotid artery which was further propelled to the middle cerebral artery, where it was left in place for 60 minutes. Following the 60-minute period of ischemia, the filament was removed and reperfusion was allowed to occur. The sham operation included the suturing of the common carotid artery for 60 minutes without filament placement. Throughout surgical procedure and during reperfusion the flow of blood was monitored using Laser Doppler Flowmetery (Moor Instruments, United Kingdom).

### Neurological deficit assessment following tMCAO surgery

All mice were scored on a 6-point neurological deficit scoring scale following sham or tMCAO procedure, and were reassessed every 24 hours for the duration of the study [10]. 0 – no neurological deficit; 1 – retracts contralateral forepaw when lifted by the tail; 2-circles to the contralateral when lifted by the tail; 3-falling to the contralateral while walking; 4 – does not walk spontaneously or is comatose; 5 – dead.

### Triphenyltetrazolium chloride (TTC) staining and quantification of infarct volume

Whole brains were removed from skull and surrounding tissue, then sectioned using a 2-mm tissue matrix. Sections were stained in TTC at 37°C for 10 minutes on each side. Sections were then imaged, and total infarct volume was measured using Image J.

### Brain tissue homogenization and single-cell isolation

Brains were harvested 24 and 72 hours following tMCAO or sham operation, and were mechanically homogenized using a razor blade. Brain homogenates were digested with collagenase D (1 mg/ml) and DNase I (200 µg/ml) for 30 minutes at 37°C. Tissues were then passed through a 100-µm cell strainer. Single cells were isolated by discontinuous Percoll gradient centrifugation.

### Lung perfusion and excision

Mice were deeply anesthetized using ketamine/xylazine combination. Once in a deep plane of anesthesia, in which mice did not respond to toe-pinch stimulus, thoracic cavity was exposed and whole-body perfusion was performed by administering 15 ml of cold phosphate-buffered saline (PBS) via the left cardiac ventricle. Lungs were excised and place into 2 ml of cold PBS supplemented with 1% FBS. Right apical lobes were weighed for homogenization.

### Lung tissue digestion and single-cell isolation

Right cardiac, diaphragmatic, azygous, and left lobes were place into digestion buffer containing collagenase D (1mg/ml) and DNase I (200 µg/ml) in Hank’s Buffered Salt Solution (HBSS). Lung lobes were inflated with 1 ml of digestion buffer and incubated at room temperature for 5 minutes. Following incubation, lung lobes were cut into pieces 2-3 mm in size and placed into 5 ml of digestion buffer. The tissues were vortexed and incubated in a 37°C water bath for 45 minutes with vigorous vortexing every 8-10 minutes. Digests were pushed through 100-µm cell strainers and contents were centrifuged for 10 minutes at 380 x g. Supernatant was discarded and tissue digests were resuspended in PBS with 1% FBS to obtain single cell suspension.

### Lung tissue homogenization

Right apical lobes were placed into a 2-ml screw cap microtube containing (3) sterile 2.3 mm zirconia/silica microbeads (BioSpec Products, Bartlesville, OK) and 200 µl of cold PBS containing 1x HALT protease inhibitor cocktail (ThermoFisher Scientific, Waltham, MA). Tissues were homogenized using the BeadBug™ benchtop microtube homogenizer on maximum speed for two minutes. Following homogenization, samples were transferred to pre-chilled tube sand centrifuged at 15870 x g for 3 minutes. Supernatants were stored at −80°C for multiplex bead array analysis.

### Lung Tissue Homogenization and Culture for the Assessment of Spontaneous Pneumonia

Mice were euthanized 24 and 72 hours following sham or tMCAO operation. Whole lungs were excised, rinsed in sterile PBS, and then mechanically homogenized in 1 ml of sterile PBS in a 7-ml glass dounce tissue grinder (Corning, Corning, NY). Tissue homogenates were passed through a 100-µm sterile cell strainer and serially diluted. Aliquots of serial dilution were plated onto Luria agar and incubated at 37°C overnight to assess for bacterial growth.

### Lung Tissue Histopathology for the Assessment of Pneumonia

Mice were euthanized 24 and 72 hours following sham or tMCAO operation. Mice were tracheally cannulated and lungs were excised. Lungs were then inflated with 10% formalin. Tissue was fixed in formalin for a minimum of 24 hours before being embedded into paraffin, sectioned, and mounted onto the slides. Sections were stained with Hematoxylin and Eosin stain (H and E) and assessed by a pathologist for the presence of histopathological features of pneumonia.

### Broncho-alveolar lavage of the Lungs

Mice were euthanized and tracheas were exposed. A cannula was inserted by a small incision into the trachea and secured with surgical suture. Thoracotomy was performed to expose lung tissue. Two fractions of a total of 3 ml cold PBS were instilled into the lungs: the first fraction of 0.4 ml was delivered, and then withdrawn following 30 seconds of continuous gentle lung massage. The second fraction of 2.6 ml were delivered in aliquots of 0.6-0.7 ml. The aliquots were delivered and withdrawn with simultaneous and continuous gentle massage of the lungs. The first fraction was centrifuged at 470 x g for 5 minutes, and supernatant was stored at −80°C for multiplex bead array analysis. The second fraction was centrifuged at 470 x g for 5 minutes, and supernatant was discarded. The cell pellets from both fractions were combined in 1 ml of cold RPMI, quantified, and analyzed by flow cytometry.

### Cell quantification and phenotyping by flow cytometry

Lung and brain single cell suspensions were quantified by the trypan blue exclusion method. Cells were blocked with anti-CD16/32 (Biolegend, San Diego, CA); immune cell types were identified using combinations of antibody listed in Table 1. All antibodies were purchased from Biolegend, San Diego, CA, except anti-Siglec F, which was purchased from BD Pharmingen, Franklin Lakes, NJ. LIVE/DEAD Fixable Dead Cell Stain was used to exclude dead cells (ThermoFisher Scientific, Waltham, MA). Samples were run on a LSRFortessa (BD Bioscineces) using FACSDiva software version 8.0, and analyzed using FlowJo version 9.9.6.

**Table 1.** Surface markers and antibody combinations for determining immune cells from the lungs and the brain following tMCAO

| Antibody | Clone | Immune Cell Type | Population | Surface Marker Expression |
| --- | --- | --- | --- | --- |
| CD45-FITC | 30-F11 | Alveolar macrophages | L1 | CD45+ Siglec F+ CD11b- |
| Siglec F-PE | E50-2440 | Eosinophils | L2 | CD45+ Siglec F+ CD11b+ |
| CD11c-Percp/Cy5.5 | N418 | CD103+ DCs | L3 | CD45+ Siglec F- CD11b- CD103+ CD11c+ MHC II+ |
| CD11b-PE/Cy7 | M1/70 | CD11b+ DCs | L4 | CD45+ Siglec F- CD11b hi CD103- CD64- MHC II+ |
| CD64-APC | X54-5/7.1 | Interstitial macrophages | L5 | CD45+ Siglec F- CD11b hi CD103- CD64+ MHC II+ |
| Live/dead-APC/Cy7 |  |  |  |  |
| CD103-BV421 | 2E7 |  |  |  |
| MHC II-BV510 | M5/114.15.2 |  |  |  |
| CD45-FITC | 30-F11 | Monocytes/moDCs | L8 | CD45+ CD11b hi Ly6C hi/int CCR2+/- Ly6G- |
| Ly6C-PE | HK1.4 | Neutrophils | L9 | CD45+ CD11b hi Ly6C int CCR2- Ly6G+ |
| CD11b-PE/Cy7 | M1/70 |  |  |  |
| Live/dead-APC/Cy7 |  |  |  |  |
| CCR2-BV421 | SA203G11 |  |  |  |
| Ly6G-BV510 | 1A8 |  |  |  |
| CD45-FITC | 30-F11 | Plasmacytoid DCs | L6 | CD45+ B220+ CD11c+ |
| CD8-PE | 53-6.7 | B cells | L7 | CD45+ B220+ CD11c- |
| NK1.1-Percp/Cy5.5 | PK136 | CD4+ T cells | L10 | CD45+ B220- CD11c- CD4+ CD8- |
| CD11c-PE/Cy7 | N418 | CD8+ T cells | L11 | CD45+ B220- CD11c- CD4- CD8+ |
| APC-B220 | RA3-6B2 | NK cells | L12 | CD45+ B220- CD11c- CD4- CD8- NK1.1+ TCRβ- |
| Live/dead-APC/Cy7 |  | NKT cells | L13 | CD45+ B220- CD11c- CD4- CD8- NK1.1+ TCRβ+ |
| CD4-BV421 | GK1.5 |  |  |  |
| TCRβ-BV510 | H57-597 |  |  |  |

### Multiplex cytokine and chemokine bead arrays

Multiplex bead arrays for cytokine and chemokine quantification were performed according to manufacturer protocol (Biolegend, catalogue #740446 and #740451). Samples were analyzed on a LSRFortessa (BD Biosciences) as described above.

### Statistical Analyses

Statistical comparison between samples was done by student’s t test. *, P < 0.05; **, P < 0.01; ***, P < 0.001. NS, not statistically different.

## Results

### Severe ischemic stroke in C57BL/6J mice does not cause spontaneous pneumonia

Several groups reported that ischemic stroke induced by tMCAO in mice causes spontaneous pneumonia, defined as lung infection without bacterial inoculation. However, previous studies have shown that the incidence of spontaneous pneumonia varies in mice with different genetic backgrounds [11], and may also depend on environmental factors such as animal housing facility conditions [12, 13]. We performed tMCAO in C57BL/6J mice by inserting a monofilament into the middle cerebral artery for 60 minutes, followed by monofilament removal to allow blood reperfusion. We obtained significant brain infarcts at 24 and 72 hours following tMCAO (Fig 1A and 1B). The percentage of infarcts in the ipsilateral cortex was over 50% and nearly 100% in the corpus striatum (Fig 1C and 1D). Correspondingly, neurological deficits of score ≥ 2 were observed (Fig 1E). In addition, we found a significant reduction in splenic cellularity (Fig 1F) in mice following tMCAO, which is a feature of severe ischemic stroke in humans and mice [3, 14]. These data indicate that induction of severe ischemic stroke was achieved. However, we could not detect spontaneous pneumonia in our mice following tMCAO. Pathological characteristics of bacterial pneumonia, such as inflammation, tissues damage, and edema were not observed in the lungs (Fig 1G and S1 Fig). The cellularity of the bronchoalveolar lavage fluid (BALF) was modestly increased in mice 24 hours following tMCAO compared to sham operation (Fig 2A). Over 90% of CD45+ cells were alveolar macrophages (Fig 2B-2E), but not neutrophils, which are commonly found in BAL during bacterial pneumonia. There was no bacterial recovery in lung homogenates 24 hours following tMCAO (S1 Table). At 72 hours, bacteria were detected in some tMCAO- and sham-operated mice (S1 Table), but the quantity was substantially lower than that which has previously been reported [3, 15]. Our findings suggest that there are factors other than the severity of the stroke that determine the incidence of spontaneous pneumonia in mice. In our studies, bacterial infection without inoculation was either absent or below detectable levels.

**Fig 1.**
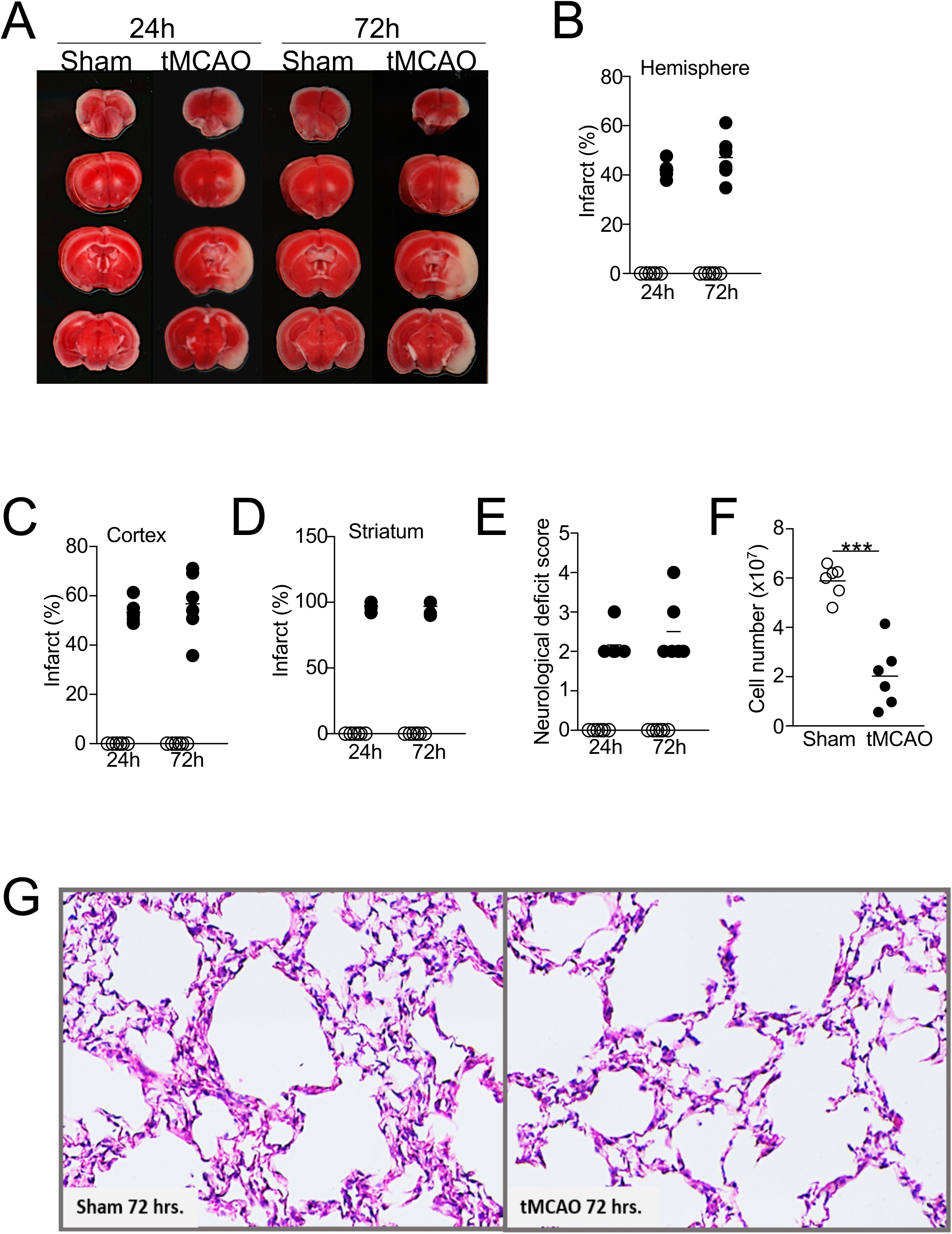
Severe ischemic stroke in C57BL/6J mice does not cause spontaneous pneumonia. Brain, spleen, and lung tissues were analyzed 24 and 72 hours following tMCAO or sham operation. (A) Representative images showing ipsilateral brain infarcts following tMCAO but not sham operation by TTC staining. (B-D) Percentage of infarcts within the ipsilateral hemisphere (B), cortex (C), and corpus striatum (D) following tMCAO (filled circle) or sham controls (open circle) quantified by Image J. (E) Neurological deficit scores of the mice 24 and 72 hours following tMCAO (filled circle) or sham controls (open circle). See Method section for score definition. (F) Cell number from the spleens of mice 72 hours following tMCAO (filled circle) or sham controls (open circle). (G) Representative images from H&E staining of lung tissues 72 hours following tMCAO (right) or sham operation (left). Images from all animals are shown in Supplemental Fig. 1. N = 6 animals per group. ***, P < 0.001.

**Fig 2.**
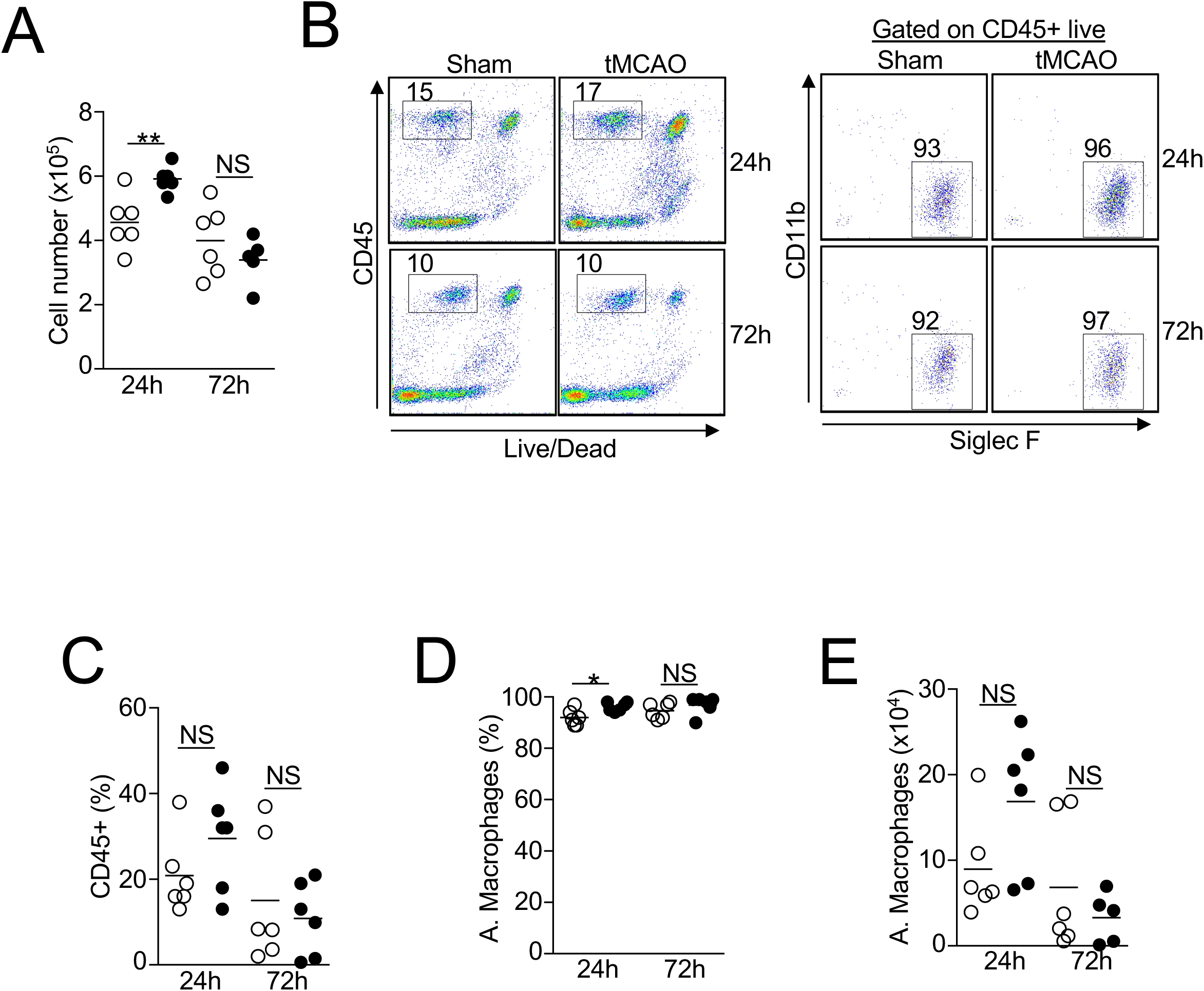
Increase in the number of alveolar macrophages in the BALF 24 hours post-ischemic stroke. (A) Total number of cells recovered from BALF 24 and 72 hours following tMCAO (filled circle) or sham operation (open circle). (B) Cellular compositions of BALF 24 and 72 hours following tMCAO. (B) Representative plots showing percentage of CD45+ cells (left) and alveolar macrophages (right), which are defined as CD45+ Siglec F+ CD11b-. (C-E) Graphs showing percentage of CD45+ cells (C); percentage (D) and number (E) of alveolar macrophages (A. Macrophages) of individual animals described in (B). tMCAO (filled circle); sham operation (open circle). N = 6 animals per group. *, P < 0.05; **, P < 0.01. NS, not statistically different.

### Alternations of the resident innate immune-cell niche in the lungs following ischemic stroke

Resident innate immune cells such as macrophages and dendritic cells (DCs) are the first line of defense during pulmonary bacterial infection. We first determined the quantitative changes of lung-resident alveolar and interstitial macrophages, CD11b+ and CD103+ conventional DCs (cDCs), plasmacytoid DCs (pDCs), and eosinophils 24 and 72 hours following tMCAO by flow cytometry (Table 1). The total number of cells and the percentage of CD45+ cells in the lungs were slightly lower but did not reach statistical significance (Fig 3A-3C). Given that some of these innate immune cells only contribute to less than 0.1% of the total lung cells, data is presented as percent change within the CD45+ population. Twenty-four hours following ischemic stroke, we found a significant increase of alveolar macrophages and CD11b+ DCs in the lungs, whereas eosinophils were reduced (Fig 3D-3G). The increase of alveolar macrophages persisted up to 72 hours following ischemic stroke, albeit to a lesser extent (Fig 3D and 3E). However, at 72 hours the number of CD11b+ DCs and eosinophils were comparable to mice with sham-operation (Fig 3D and 3F). The percentages of interstitial macrophages, CD103+ DCs, and pDCs were unchanged at both time points (Fig 3D, 3H-3K). These data suggest that ischemic stroke alters specific subsets of immune cells in the pulmonary environment, particularly 24 hours post-stroke.

**Fig 3.**
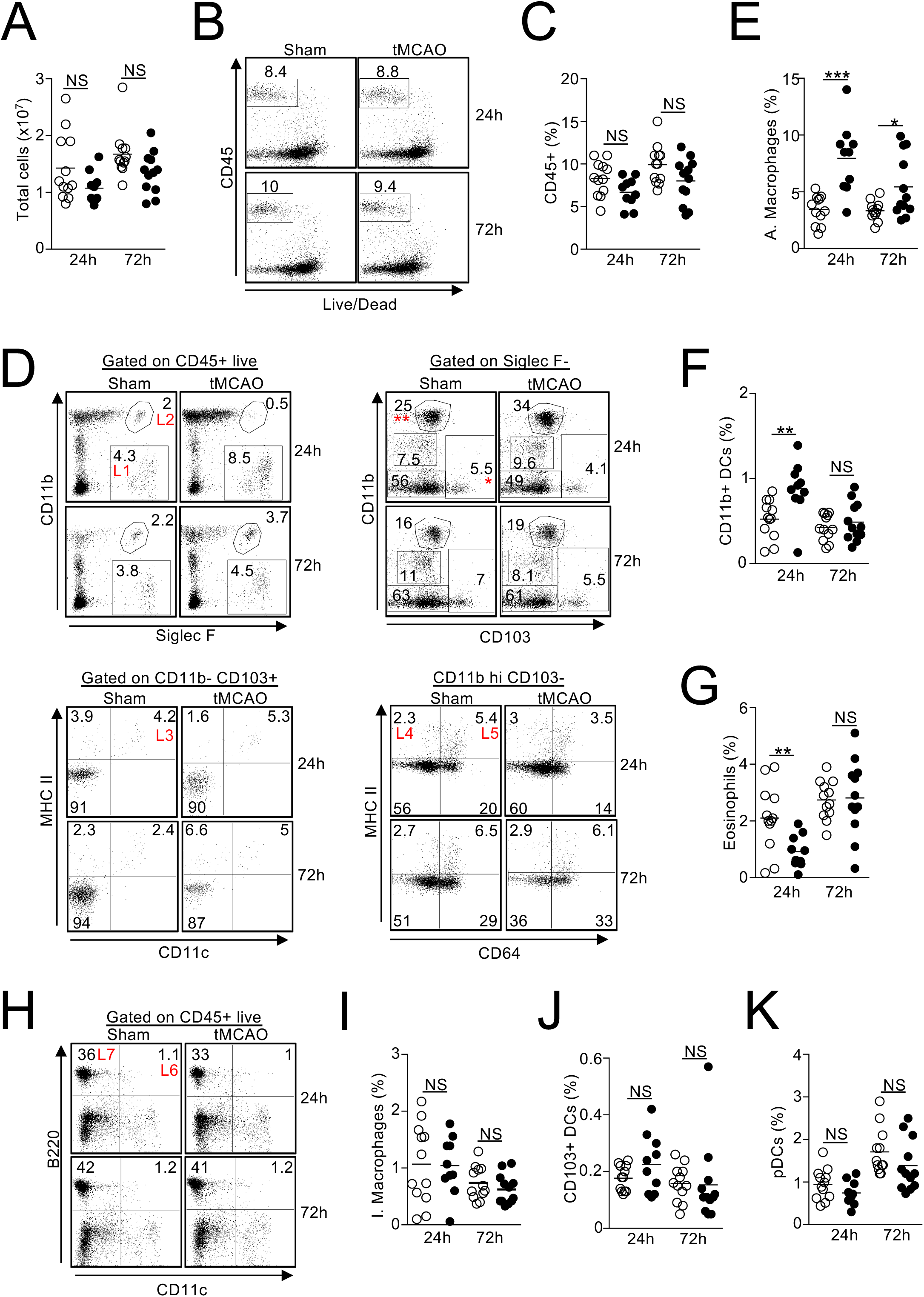
Alterations in the resident innate immune cell niche in the lungs following ischemic stroke. Lung tissues were excised 24 and 72 hours following tMCAO (filled circle) or sham operation (open circle), resident innate immune cells in the lungs were analyzed by flow cytometry, defined by surface markers listed on Table 1. (A) Graph showing total number of cells in the lungs of individual animals. (B) Representative plots showing percentage of CD45+ cells. (C) Graph showing percentage of CD45+ cells in the lungs of individual animals. (D) Representative plots showing the identification of alveolar macrophages (top left, L1), eosinophils (top left, L2), CD103+ DCs (bottom left, L3), CD11b+ DCs (bottom right, L4), and interstitial macrophages (bottom right, L5). CD103+ DCs were first gated on CD103+ CD11b-cells (top right, *). CD11b+ DCs and interstitial macrophages were first gated on CD11b hi CD103-cells (top right, **). (E-G) Graphs showing percentage of alveolar macrophages (E), CD11b+ DCs (F), and eosinophils (G) of individual animals. (H) Representative plots showing that identification of pDCs (L6), and B cells (L7, discussed in Fig. 5). (I-K) Graphs showing percentage of interstitial macrophages (I), CD103+ DCs (J), and pDCs (K) of individual animals. N = 12 animals per group. **, P < 0.01; ***, P < 0.001. NS, not statistically different.

### Increased infiltration of neutrophils but not monocytes to the lungs following ischemic stroke despite an elevation of CCL2

In addition to the activation of resident innate immune cells, neutrophil and monocyte infiltration to the lungs plays a critical role in bacterial clearance [16–18]. We determined whether ischemic stroke affects trafficking of these cells to the lungs. Flow cytometric analysis revealed an increase of neutrophils in the lungs 24 and 72 hours following tMCAO (Fig 4A and 4B), whereas the percentage of monocytes and monocyte-derived dendritic cells (moDCs) was not changed (Fig 4A and 4C), despite an increase in the monocyte chemoattractant CCL2 (Fig 4D). Interestingly, there was a massive infiltration of monocytes and moDCs to the brain following ischemic stroke (Fig 4E and 4F), however, the number of neutrophils in the brain was unchanged (Fig 4E and 4G). These data suggest that ischemic stroke differentially directs monocyte and neutrophil trafficking into the brain and lungs respectively, which may contribute to the post-stroke immunosuppression.

**Fig 4.**
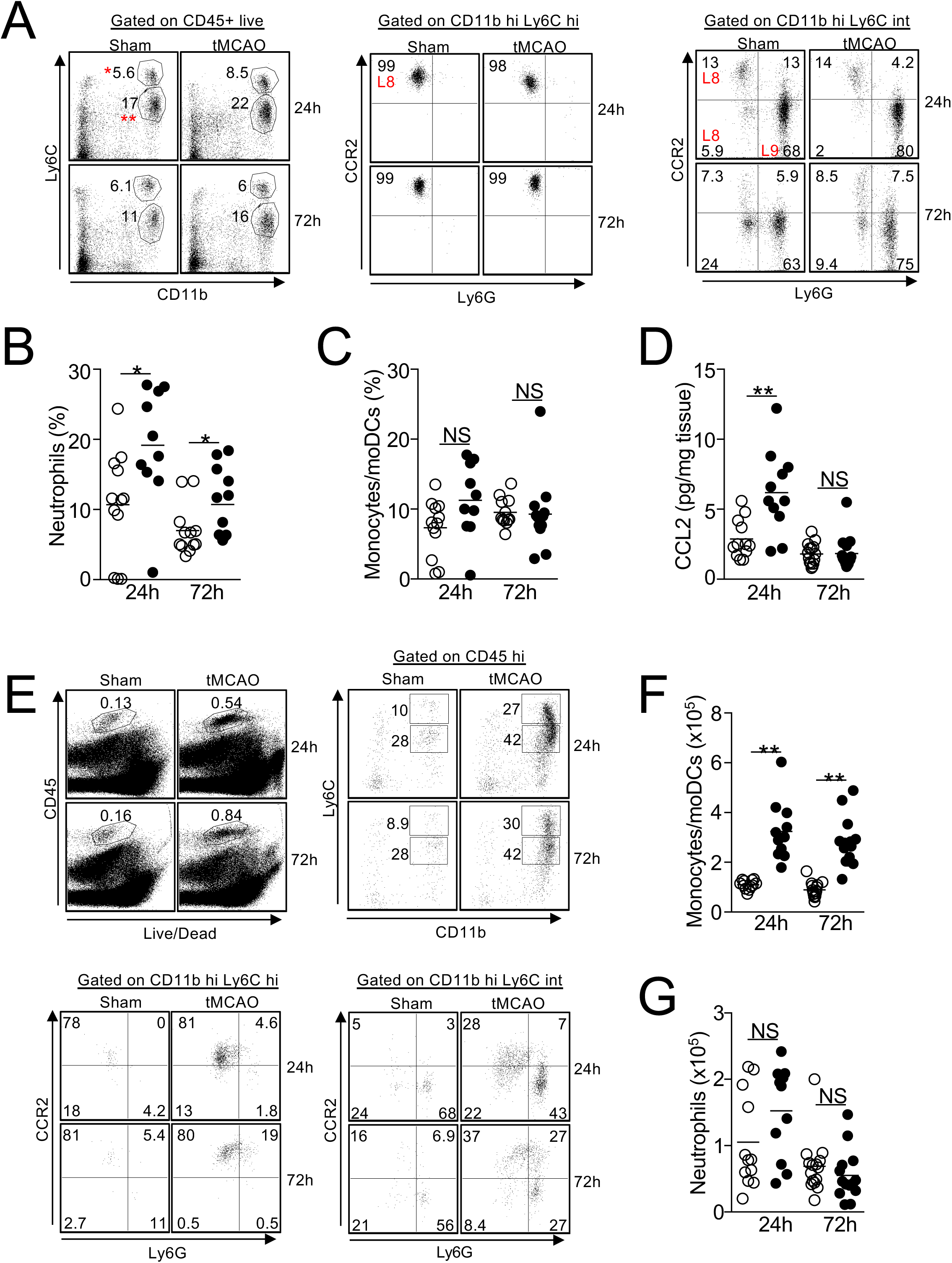
Increased infiltration of neutrophils but not monocytes to the lungs following ischemic stroke despite an elevation of CCL2. (A-C) Lung tissues were excised 24 and 72 hours following tMCAO or sham operation. Monocytes/moDCs and neutrophils in the lungs were analyzed by flow cytometry, defined by surface markers listed on Table 1. (A) Representative plots showing the identification of monocytes/moDCs (L8) and neutrophils (L9) in the lungs. Monocytes/moDCs were defined as CD45+ Ly6C hi (*) CCR2 hi (middle) and Ly6C intermediate (**) CCR2+/- (right). Neutrophils were defined as CD45+ Ly6C intermediate (**) Ly6G+ (right). (B-C) Graphs showing percentage of neutrophils (B) and monocytes/moDCs (C) of individual animals 24 and 72 hours following tMCAO (filled circle) or sham operation (open circle). (D) Lung tissues were homogenized 24 and 72 hours following tMCAO (filled circle) or sham operation (open circle), level of CCL2 was determined by multiplex bead array. (E-G) Brain tissues were excised 24 and 72 hours following tMCAO or sham operation, monocytes/moDCs and neutrophils in the brains were analyzed by flow cytometry. (E) Representative plots showing the identification of monocytes/moDCs and neutrophils in the brains, which were defined as in (A). (F-G) Graphs showing number of monocytes/moDCs (F) and neutrophils (G) of individual animals 24 and 72 hours following tMCAO (filled circle) or sham operation (open circle). N = 12 animals per group. *, P < 0.05; **, P < 0.01. NS, not statistically different.

### Ischemic stroke leads to a significant loss of lymphocytes in the lungs independent of apoptosis

Previous studies have shown that ischemic stroke leads to the apoptosis of T cells, B cells, and NK cells in the spleen, blood, and thymus [3]. However, how ischemic stroke alters the lymphocyte population in the lungs is not clear. We found that the percentage of CD4+ T cells, CD8+ T cells, and B cells were significantly reduced 24 and 72 hours following tMCAO (Figs 3H, 5A-5D). NK cells were reduced at 24 but not 72 hours following tMCAO (Fig 5A and 5E), whereas NKT cells were unaltered in both time points (Fig 5A and 5F). We speculated that the loss of lymphocytes in the lungs was caused by increased apoptosis as was previously described in other tissues. We determined the percentage of apoptotic cells by annexin V staining. The percentage of apoptotic CD4+ T cells, CD8+ T cells, B cells, and NK cells were comparable in sham-operated and tMCAO-induced mice (Fig 5G-5J), suggesting that unlike other tissues, ischemic stroke leads to a significant loss of lymphocytes in the lungs but it is unrelated to the induction of apoptosis. We then investigated whether the reduction of lymphocytes in the lungs was caused by cell trafficking to the brain after ischemic stroke. Surprisingly, the number of CD4+ T cells, CD8+ T cells, B cells, and NK cells in the brain was either unchanged or reduced post-stroke, indicating that the loss of lymphocytes in the lungs following tMCAO likely does not the result from cell infiltration to the brain (Fig 6A-6E).

**Fig 5.**
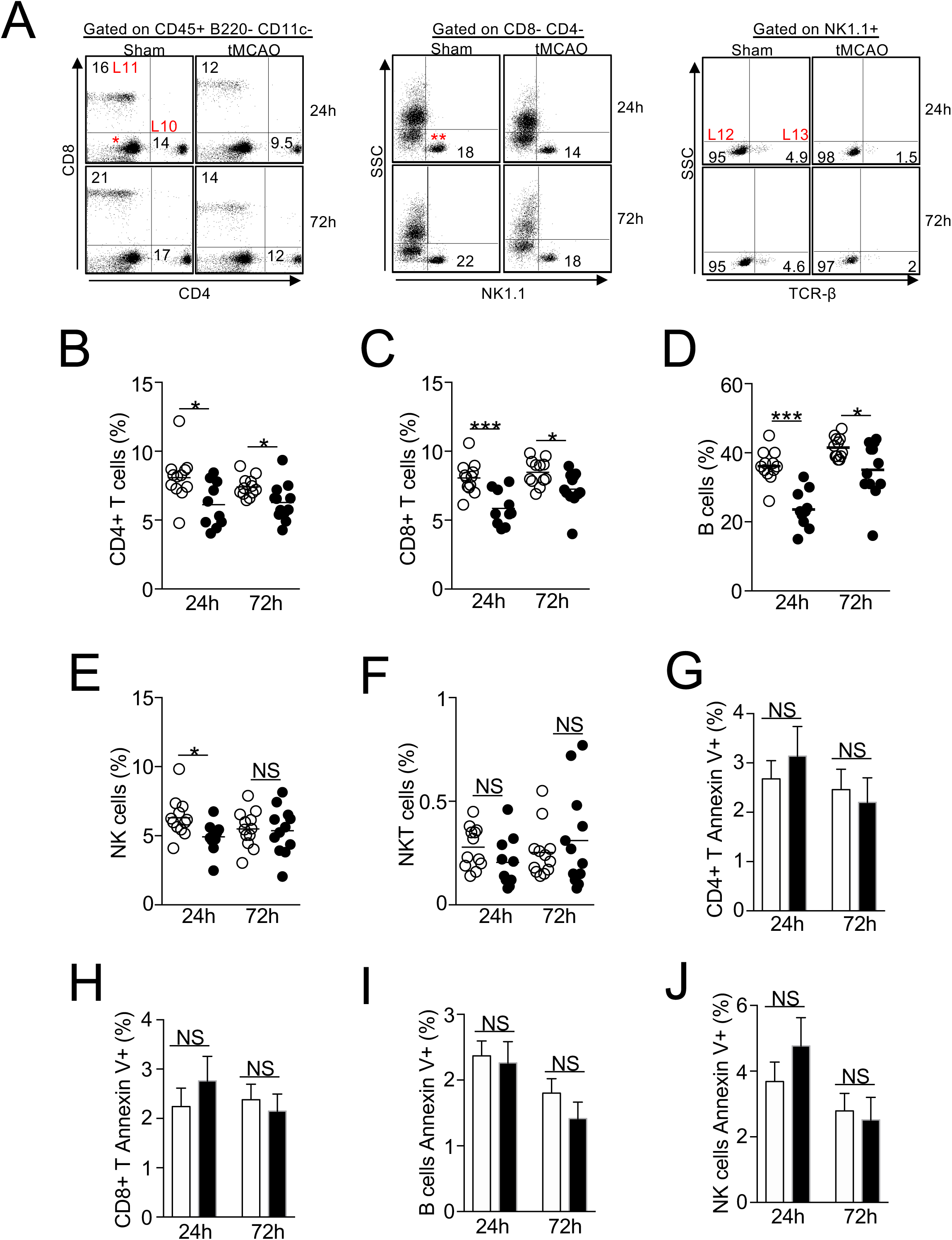
Ischemic stroke leads to a significant loss of lymphocytes in the lungs independent of apoptosis. (A) Representative plots showing the identification of lymphocytes in the lungs by surface markers listed on Table 1. CD4+ (L10) and CD8+ (L11) T cells were first gated on CD45+ B220- CD11c- cells shown in Fig. 3H. NK cells (L12) and NKT cells (L13) were gated on CD4- CD8- cells (*), then further gated on NK1.1+ cells (**). Representative plots for the identification of B cells shown in Fig. 3H. (B-F) Graphs showing percentage of CD4+ T cells (B), CD8+ T cells (C), B cells (D), NK cells (E), and NKT cells (F) of individual animals 24 and 72 hours following tMCAO (filled circle) or sham operation (open circle). (G-H) Graphs showing percentage of annexin V+ CD4+ T cells (G), CD8+ T cells (H), B cells (I), and NK cells (J) 24 and 72 hours following tMCAO (filled bar) or sham operation (open bar). N = 12 animals per group. *, P < 0.05; ***, P < 0.001. NS, not statistically different.

**Fig 6.**
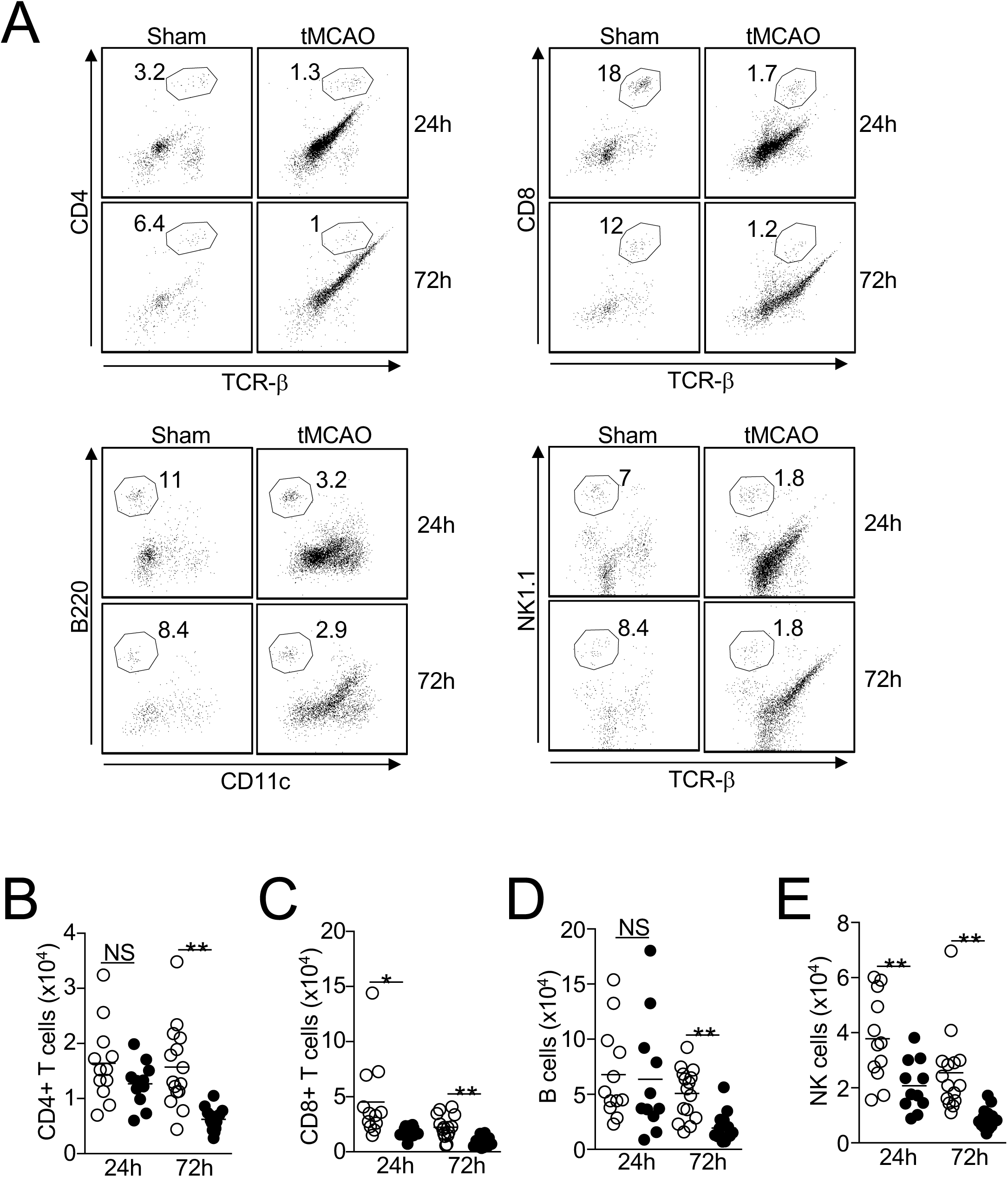
Loss of lymphocytes in the lungs following tMCAO is not the result of cell migration to the brain. (A) Representative plots showing the identification of CD4+ T cells (CD4+ TCR-β+, top left); CD8+ T cells (CD8+ TCR-β+, top right); B cells (B220+ CD11c-, bottom left); and NK cells (NK1.1+ TCR-β-, bottom right) in the brains. Cells were first gated on CD45 hi cells shown in Fig. 4E. (B-E) Graph showing number of CD4+ T cells (B), CD8+ T cells (C), B cells (D), and NK cells (E) of individual animals 24 and 72 hours following tMCAO (filled circle) or sham operation (open circle). N = 12 animals per group. *, P < 0.05; **, P < 0.01. NS, not statistically different.

### Ischemic stroke suppresses the production of multiple chemokines and cytokines in the lungs

Chemokine and cytokine expression play a key role in regulating immune-cell trafficking and function. Using multiplex bead arrays, we measured the expression of a total of 25 chemokines and cytokines in the lungs following ischemic stroke that are known to control inflammatory responses. Among the 13 chemokines we detected, the levels of CCL3, CCL5, CCL22, CXCL5, CXCL9, and CXCL10 were significantly reduced at both 24- and 72-hour time points following tMCAO (Fig 7A-7F). The levels of CCL4, CCL17, and CCL20 were reduced only at one time point (Fig 7G-7I). The suppression of chemokine production was not caused by a general lung dysfunction as the levels of CCL11, CXCL1, and CXCL13 were comparable between the sham-operated and tMCAO mice (Fig 7J-7L). Interestingly, CCL2 was the only chemokine measured that was increased in the lungs following tMCAO (Fig 4D). In contrast to the lung tissues, chemokine levels in the BALF were mostly unchanged following tMCAO, except for a reduction in CCL22 at both time points and CCL20 at 72 hours post-stroke (S2 Fig). Suppression of chemokine expression in the lungs may explain the loss of lung lymphocytes post-ischemic stroke.

**Fig 7.**
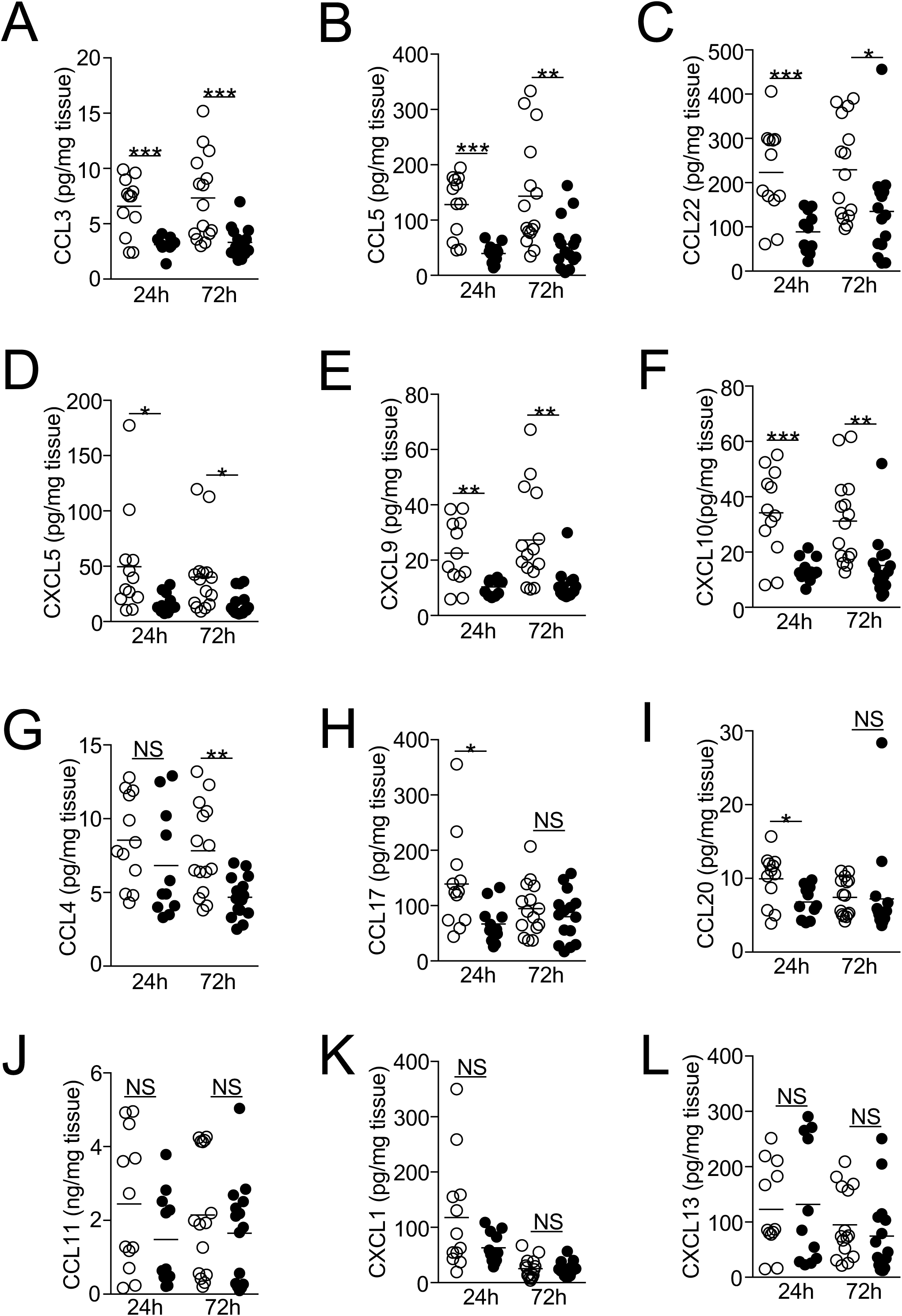
Ischemic stroke suppresses the production of multiple chemokines in the lungs. (A-L) Lung tissues were homogenized 24 and 72 hours following tMCAO (filled circle) or sham operation (open circle), level of CCL3 (A), CCL5 (B), CCL22 (C), CXCL5 (D), CXCL9 (E), CXCL10 (F), CCL4 (G), CCL17 (H), CCL20 (I), CCL11 (J), CXCL1 (K), CXCL13 (L) in the lungs of individual animals was determined by multiplex bead array. N = 12-15 animals per group. *, P < 0.05; **, P < 0.01; ***, P < 0.001. NS, not statistically different.

Proinflammatory cytokines such as IL-1β, TNF-α and IL-6 are critical for promoting bacterial clearance during lung infections [19, 20]. IFN-γ and IL-17A are signature cytokines of Th1 and Th17 cells, both of which support innate-cell activation and migration [21, 22]. We found that the levels of IL-1β, TNF-α, IFN-γ, IL-17A, and IL-27 were reduced 72 hours following ischemic stroke (Fig 8A-8E). Importantly, we found that IL-1α was abundantly expressed in the lungs of the sham-operated mice relative to other cytokines we measured (∼50 pg/mg of tissue), but its level was significantly reduced at both 24- and 72-hour time points post-ischemic stroke (Fig 8F). Levels of IL-6, IL-12p70, IL-23, IL-10, IFN-β, and GM-CSF (Fig 8G-8L) were unaltered following tMCAO. Significant changes in cytokine levels in BALF following tMCAO were not observed (S3 Fig). Overall, these data suggest that ischemic stroke creates an immunosuppressive milieu in the lungs by decreasing the production of multiple proinflammatory chemokines and cytokines, which may be responsible for the higher susceptibility to bacterial infection following ischemic stroke.

**Fig 8.**
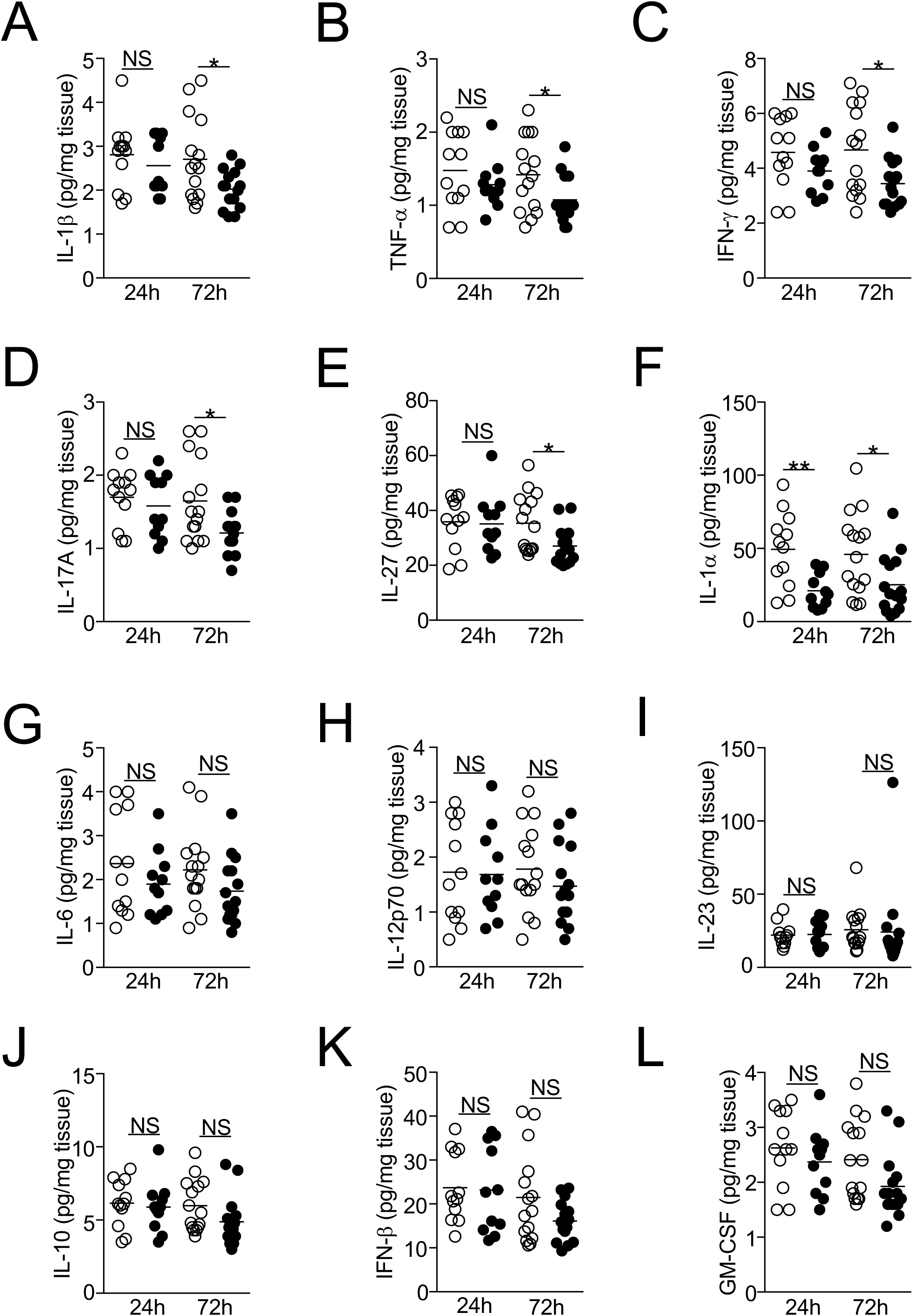
Ischemic stroke suppresses the production of multiple cytokines in the lung. (A-L) Lung tissues were homogenized 24 and 72 hours following tMCAO (filled circle) or sham operation (open circle), level of IL-1β (A), TNF-α (B), IFN-γ (C), IL-17A (D), IL-27 (E), IL-1α (F), IL-6 (G), IL-12p70 (H), IL-23 (I), IL-10 (J), IFN-β (K), GM-CSF (L) in the lungs of individual animals was determined by multiplex bead array. N = 12-15 animals per group. *, P < 0.05; **, P < 0.01. NS, not statistically different.

## Discussion

Our study demonstrated that ischemic stroke directly impacts the immune-cell niche, and the availability of multiple chemokines and cytokines in the lungs, which was previously unexplored. These changes were not the result of nor the cause of spontaneous pneumonia as reported in other studies [3, 15]. Consistent with our observations, recent studies have also shown that severe ischemic stroke in mice does not cause spontaneous pneumonia, but rather those mice are highly susceptible to bacterial infection induced by intra-tracheal inoculation [23]. Using tMCAO model, Stanley et al showed that SAP is caused by the translocation and dissemination of commensal bacteria from the intestinal tracts to lungs, and SAP does not occur in mice housed in a germ-free environment [15]. These observations strongly suggest that the incidence of SAP is determined by the composition of gut microbiota, and this speculation is further supported by the fact that mice with different genetic backgrounds develop SAP with various severity [11]. Here we propose that the incidence of SAP may also depend on factors other than genetic backgrounds such as the source of the animals and hygiene condition of the housing facility, which is commonly observed in many animal disease models [24]. Importantly, it is still unclear whether gut bacterial translocation to the lungs is the cause of clinical SAP. While some SAP-causative pathogens such as *Klebsiella pneumoniae* and *Escherichia coli* are commonly found in the gut microflora, *Staphylococcus aureus*, *Pseudomonas aeruginosa*, and *Streptococcus pneumoniae* are not [25]. It is possible that clinical SAP results from a combination of nosocomial infection and bacterial translocation from the gut microflora. This possibility can be explored by comparing the composition of the gut microflora of patients with severe ischemic stroke that develop SAP with those patients that do not develop SAP. In any case, stroke-induced immunosuppression is a major cause of SAP in clinical and animal studies.

Currently, it is still unclear how ischemic stroke-mediated immune-cell dysfunctions in remote tissues such as the spleen and the thymus lead to pneumonia. Our data suggested that alternations in the pulmonary immune-cell niche following ischemic stroke may at least partly explain the lung-specific immunodeficiency that may lead to a higher susceptibility to bacterial infections. A recent study has shown that neutrophils from the mouse bone marrow have an impaired chemotactic ability post-stroke in an *in vitro* model [9]. However, neutrophil infiltration to the lungs was observed in both spontaneous- or aspiration-induced pneumonia following tMCAO [3, 5]. We observed an increase in alveolar macrophages and the infiltration of neutrophils in the lungs following tMCAO, but their bacterial clearance capability remains to be elucidated. Oxidative burst and NETosis were reduced in neutrophils isolated from patients who experienced ischemic stroke [26]. To connect these functional deficits with SAP, we are currently investigating the anti-bacterial activity of alveolar macrophages and neutrophils in the lungs following ischemic stroke using reporter mouse strains coupled with live-imaging techniques. In addition, recruitment of monocytes during lung infection is critical for bacterial clearance [27]. Clinically, reduction of HLA-DR expression in monocytes is a prognostic marker of stroke-induced immune suppression and SAP [2, 28, 29]. Interestingly, we found that ischemic stroke increases the expression of the monocyte chemoattractant CCL2 in the lungs, yet monocyte infiltration into the lungs was not observed. Conversely, following ischemic stroke, we observed a massive infiltration of monocytes into the brain, which may account for temporal peripheral immune exhaustion following ischemic stroke, leading to a higher susceptibility to bacterial infections.

This study is the first report showing a significant reduction of lymphocytes in the lungs following ischemic stroke that was not caused by the induction of cell death, but was associated with the decreased production of multiple chemokines. The effect of ischemic stroke on regulating chemokine expression in the lungs is not known. Our data suggest that suppression of chemokine expression may “pre-condition” the lungs to become vulnerable to bacterial infections. Particularly, CCL5 and CCL22 were abundantly expressed in the sham-operated mice (> 100 pg/mg of tissue), but their levels were significantly reduced following tMCAO. CCL5 is a potent chemoattractant of T cells to the site of inflammation, which may explain the reduction of these cells in the lungs following tMCAO. CCL5 antibody treatment in mice challenged with *Streptococcus pneumoniae* caused reductions in CD4^+^ and CD8^+^ T lymphocytes, resulting in dysregulation of a critical phase of the adaptive response. The loss of CCL5 ultimately led to a switch in bacteria from carrier state to lethal state of infection [30]. CCL22 is known to promote Th2-mediated immune responses such as airway hypersensitivity, atopic dermatitis, and eosinophilic pneumonia [31–33], but its role in bacterial infections is unclear. CCL22 is mainly produced by macrophages and dendritic cells [33, 34]; reduction of CCL22 in the lungs indicates a possible functional impairment of these cells following ischemic stroke, which warrants further investigation.

There is an urgent need to develop effective immunotherapeutic strategies for SAP. Elucidating dysregulations in the pulmonary immune-cell niche and the functions of the cells within this niche is a critical first step to achieve this goal. We demonstrated that ischemic stroke directly impacts pulmonary immunity. Restoration of the chemokine availability in the lungs may potentially prevent SAP in stroke patients.

## Acknowledgments

This work was supported by NIH grant P20 GM109098 to Edwin Wan and Xuefang Ren, NIH grant 5U54GM104942 and the American Heart Association Scientist Development Grant 16SDG31170008 to Xuefang Ren. Flow Cytometry experiments were performed in the WVU Flow Cytometry & Single Cell Core Facility, which is supported by NIH grants S10OD016165, U57GM104942, P30GM103488, and P20GM103434. Imaging experiments were performed in the WVU Imaging Facilities, which is supported by the WVU Cancer Institute, the WVU HSC Office of Research and Graduate Education, and NIH grants P20RR016440, P30GM103488, P20GM121322, U54GM104942, P20GM103434, and P20GM103434. We thank Sarah Milne for technical support.

## Authorship

E.W. and B.F. designed and performed experiments, analyzed data, and wrote the paper. K.M., C.A., and W.Z. performed experiments and analyzed data. H.H. and X.R. performed experiments. J.C. analyzed data. E.W. supervised the project.

---

**S1 Fig.**
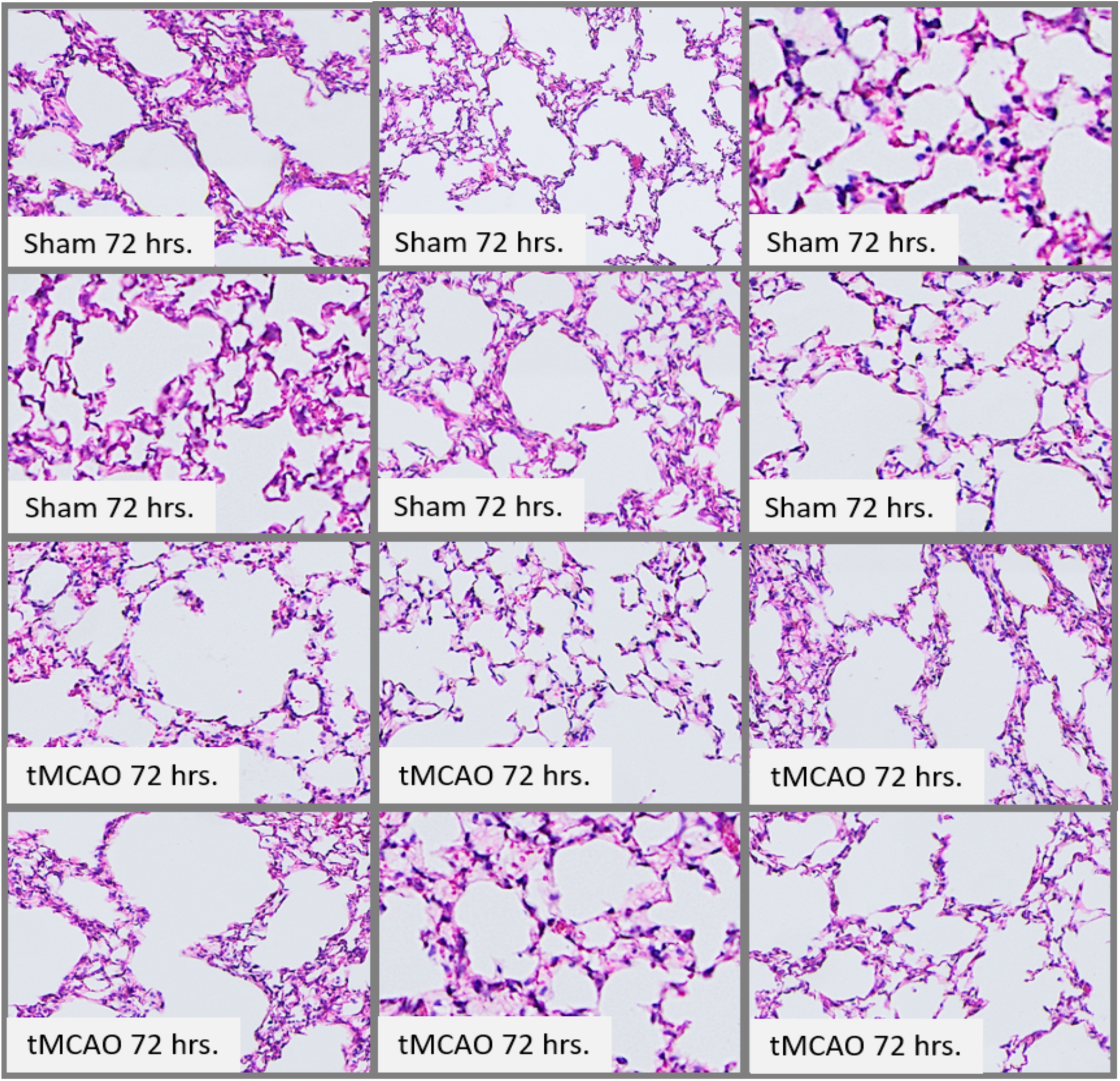
Severe ischemic stroke in C57BL/6J mice does not cause spontaneous pneumonia. Lung pathology was examined 72 hours following tMCAO (bottom) or Sham operation (top). Shown are H and E images from all animals examined. N = 6 animals per group.

**S2 Fig.**
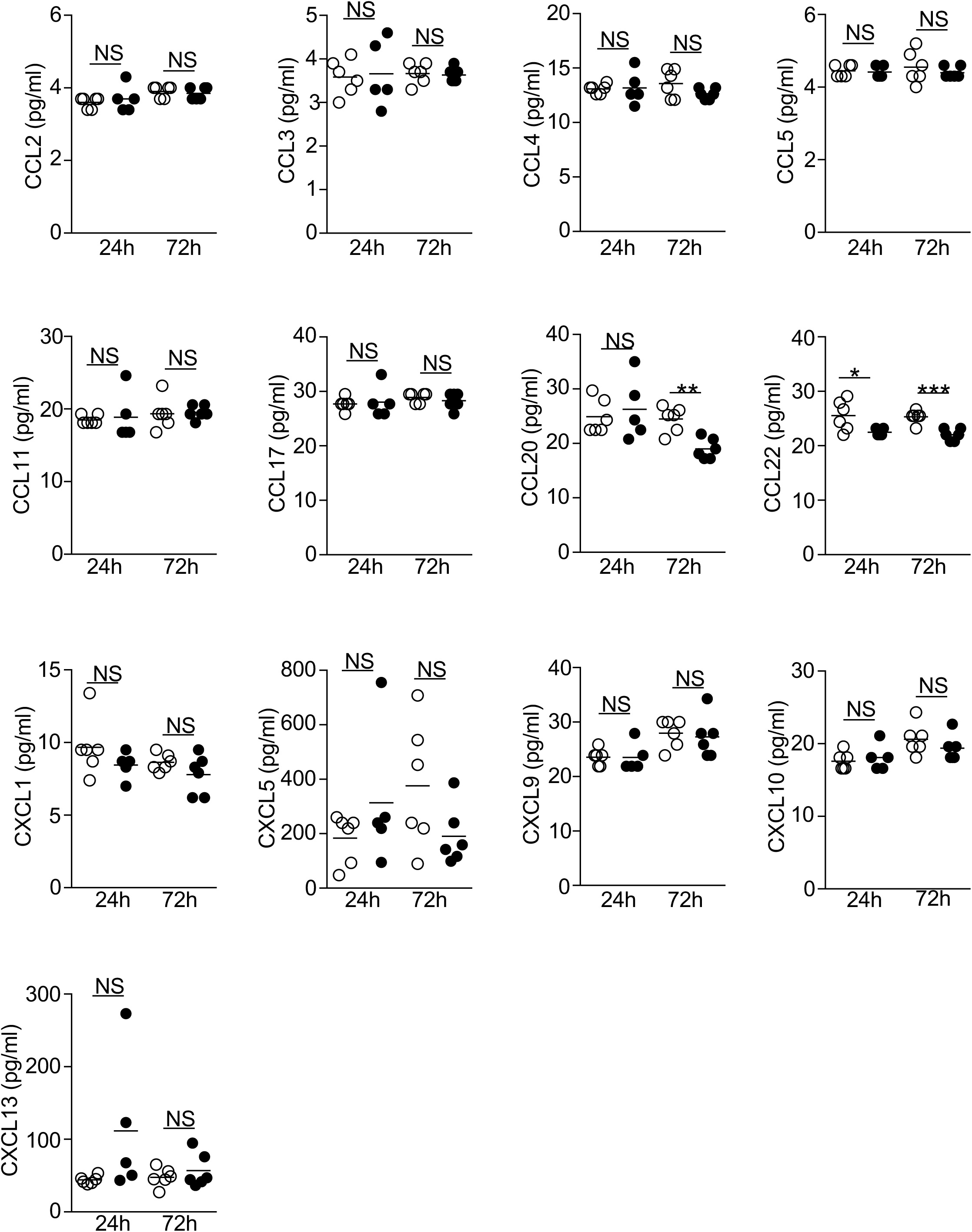
Ischemic stroke reduces the level of CCL20 and CCL22 in the BALF. BALF was collected 24 and 72 hours following tMCAO (filled circle) or sham operation (open circle), level of chemokines described in Fig. 7 was determined. N = 5-6 per group. *, P < 0.05; **, P < 0.01; ***, P < 0.001. NS, not statistically different.

**S3 Fig.**
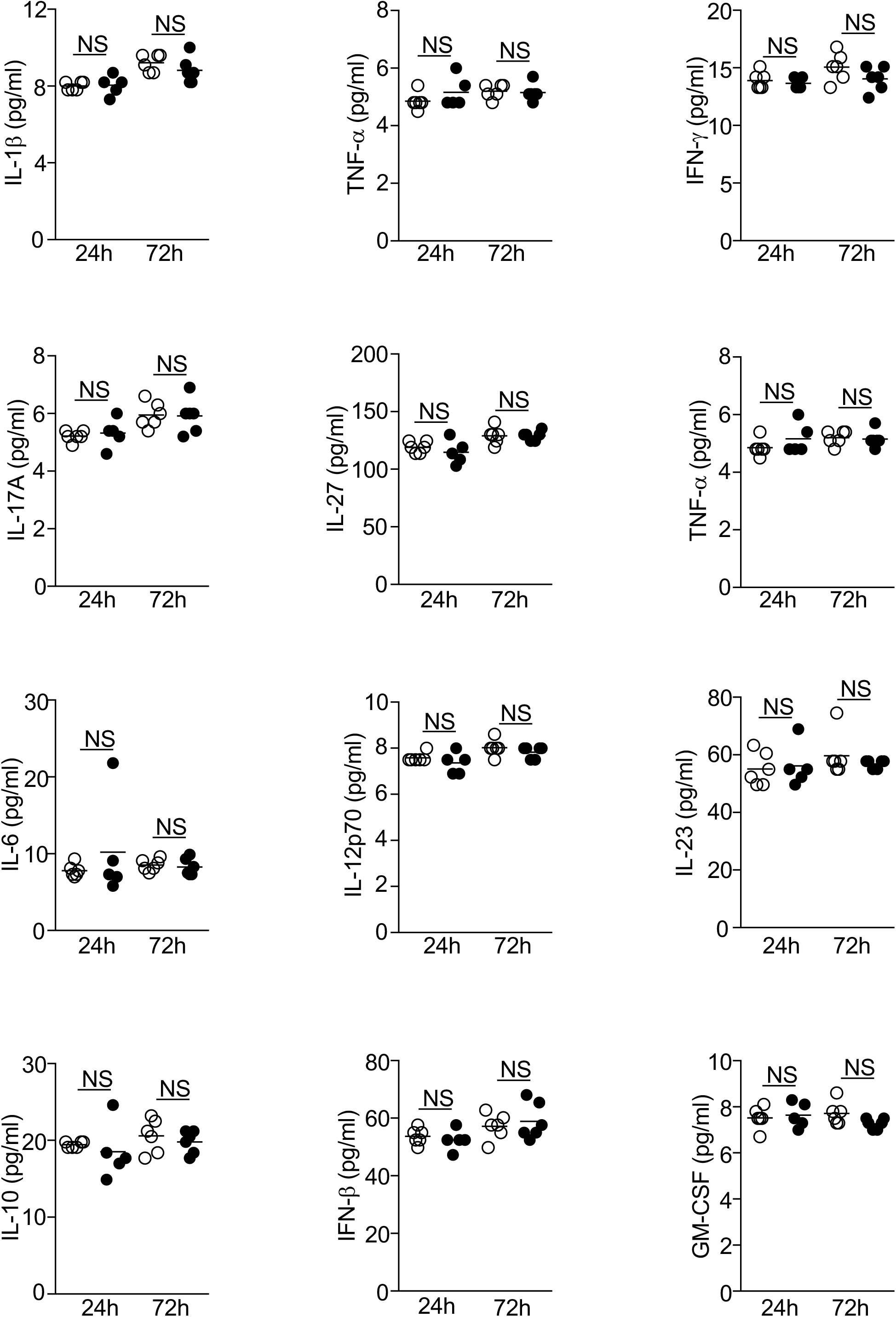
Ischemic stroke does not change the level of proinflammatory cytokines in the BALF. BALF was collected 24 and 72 hours following tMCAO (filled circle) or sham operation (open circle), level of cytokines described in Fig. 8 was determined. N = 5-6 per group. NS, not statistically different.

**Supplemental Table 1.** Recovery of bacteria from the lungs 24 or 72 hours following.

| **Surgery group** | **Hours Post-Surgery** | **Neurological Score** | **Total CFU Recovered Per Lung** |
| --- | --- | --- | --- |
| sham | 24 | 0 | 0 |
| sham | 24 | 0 | 0 |
| sham | 24 | 0 | 0 |
| tMCAO | 24 | 2 | 0 |
| tMCAO | 24 | 2 | 0 |
| tMCAO | 24 | 2 | 0 |
| tMCAO | 24 | 2 | 0 |
| sham | 72 | 0 | 0 |
| sham | 72 | 0 | 0 |
| sham | 72 | 0 | 0 |
| sham | 72 | 0 | 2575 |
| sham | 72 | 0 | 0 |
| sham | 72 | 0 | 3875 |
| tMCAO | 72 | 2 | 0 |
| tMCAO | 72 | 2 | 0 |
| tMCAO | 72 | 2 | 6000 |
| tMCAO | 72 | 2 | 2100 |
| tMCAO | 72 | 2 | 2325 |
| tMCAO | 72 | 2 | 0 |
| tMCAO | 72 | 2 | 800 |

